# MetaBeeAI: an AI pipeline for structured evidence extraction from biological literature

**DOI:** 10.1101/2025.11.24.690154

**Authors:** Rachel H. Parkinson, Henry Cerbone, Mikael Mieskolainen, Shuxiang Cao, Alasdair D. Wilson, Sergio Albacete, Emily B. Armstrong, Chris Bass, Cristina Botías, Andrew Brown, Angela J. Hayward, Lina Herbertsson, Andrew K. Jones, Nicolas Nagloo, Elizabeth Nicholls, Elisa Rigosi, Fabio Sgolastra, Harry Siviter, Dara A. Stanley, Lars Straub, Edward A. Straw, Rafaela Tadei, Kieran Walter, Heloise F. Stevance, Ryan K. Daniels, Ben Lambert, Stephen Roberts

## Abstract

The volume and complexity of scientific literature are expanding rapidly, making it increasingly difficult to extract and synthesise information across studies. This challenge is particularly acute in the biological sciences, where evidence spans multiple levels of organisation and heterogeneous experimental designs. Large Language Model (LLM) pipelines offer a scalable route to evidence synthesis, but many existing approaches lack transparency, modularity, and effective mechanisms for human oversight. We present MetaBeeAI, an open-source, modular pipeline that integrates established LLM techniques into a coherent, auditable workflow for structured data extraction in biology. MetaBeeAI combines modular prompting, multi-pass extraction, and expert-in-the-loop validation within an interface that presents model outputs alongside source text, enabling inspection, correction, and iterative refinement. The pipeline produces machine-readable records of prompts, configurations, and expert annotations, supporting reproducibility and continuous improvement. We apply MetaBeeAI to 924 research papers on bees and pesticides, extracting structured information on species, compounds, exposure designs, and experimental context. Evaluation demonstrates improved consistency, convergence with expert judgement, and robustness across heterogeneous biological studies, highlighting the value of expert-guided refinement. MetaBeeAI provides a transparent and extensible framework for scalable evidence synthesis, supporting reliable integration of LLMs into biological research workflows.

## 1. Introduction

The volume of bioscience literature has expanded exponentially in the past decade, outpacing researchers’ ability to keep abreast of current information. This surge has been accompanied by increasing complexity, fractured data standards, and increasingly specialized nomenclature and practices. As a result, conducting systematic literature reviews, a cornerstone of evidence synthesis and policy development, has become increasingly challenging. This is especially clear for experiments in data-rich fields that report effects and interactions across hierarchical levels, such as in ecology (Cavigelli et al., 2021; Köhler and Triebskorn, 2013).

The explosion of large language models (LLMs) has been proposed as a potential solution to this information bottleneck, yet many AI-based tools fall short (Channing and Ghosh, 2025). While these tools are useful, they often fail to meet the levels of transparency and adaptability required to be applied to arbitrary subfields in the biosciences. Additionally, it requires great technical expertise to fine-tune existing models to perform bespoke scientific literature review tasks, especially in fields that differ from those standardly available in the training sets. Most pretrained models are biased toward computer science and machine learning corpora, limiting performance in underrepresented domains such as ecology (Dorm et al., 2025).

To address these limitations, we developed MetaBeeAI, a modular LLM-based pipeline for structured evidence extraction with expert-in-the-loop validation. Users can define a literature corpus, design prompt templates, and iteratively assess model outputs against the source text. The modular architecture keeps MetaBeeAI adaptable as LLM capabilities and tooling evolve, while built-in transparency, testing, and continuous integration support reproducibility, long-term sustainability, and the creation of rich datasets for reinforcement learning from human feedback (Ma et al., 2024; Ziegler et al., 2020).

We demonstrate MetaBeeAI in bee ecology and ecotoxiology. This field, while having great real-world importance for food security and ecosystem functioning (Klein et al., 2006; Potts, 2016;, EFSA), is not well represented in current literature and benchmark datasets used to fine-tune LLMs (Cui et al., 2025). Bees are critical for global food production and biodiversity (Patel et al., 2021; Parreño et al., 2022), yet their populations are threatened by agricultural pesticides and interacting stressors (Goulson et al., 2015). Research on pesticide impacts spans molecular to ecosystem levels, generating an ever-growing and highly heterogenous literature base that is increasingly difficult to synthesise manually. Through a series of workshops, we curated and validated 924 research articles to evaluate MetaBeeAI’s performance. Here, we present the outcomes of that validation, assess the pipeline’s reliability and generalizability, and provide all associated data and code to facilitate adoption of MetaBeeAI for systematic literature review in other scientific domains.

## 2. Evaluation of existing methods

The rapid expansion of large language model (LLM)–based tools for scientific text analysis has generated a diverse ecosystem of literature-review pipelines, ranging from domain-specific assistants to general-purpose research agents. However, despite significant progress, most current systems remain limited in scalability, transparency, or adaptability when applied to specialized scientific domains such as biology and ecology. Many popular tools emphasize document summarization or citation tracking rather than full-text data extraction and structured synthesis. For example, current versions of ChatGPT Plus, Edu, and Pro support only 20–40 uploaded files per project for summarization and synthesis tasks (OpenAI, 2025), whereas our pipeline has been tested on datasets of around 1,000 papers and can scale well beyond that. This technical ceiling fundamentally constrains the feasibility of comprehensive evidence synthesis at the scale required for systematic reviews in data-rich disciplines.

A review of existing LLM-based literature platforms (Table A.3) reveals that, while several tools can perform partial or modular review functions, none offer the combined attributes of full-text ingestion, modular multi-stage processing, expertin-the-loop validation, and benchmarking that define MetaBeeAI. Systems such as Elicit (Elicit, 2025), PaperQA2 (Lála et al., 2023), AstaAI (Allen Institute for AI, 2024), and SciSpace (Inc., 2025) allow users to query scientific corpora and retrieve summaries but lack structured data output or fine-grained traceability to source text. Others, including Consensus (Consensus, 2024) and Perplexity (Deep Research) (Inc, 2025) emphasize answer generation or conversational exploration rather than standardized, reproducible synthesis workflows.

MetaBeeAI differs fundamentally in that it operates as a multi-pass, modular LLM pipeline capable of parsing, extracting, and benchmarking structured information from full-text PDFs. Unlike single-prompt summarization systems, MetaBeeAI implements explicit stages for chunk-level relevance filtering, iterative prompt refinement, and human expert validation. Each step produces transparent, auditable outputs that can be inspected or replaced without retraining the underlying model. This modular architecture allows continual improvement as newer LLMs and auxiliary tools emerge, providing a long-lived framework for evidence synthesis.

Table A.3 summarizes the capabilities of the principal literature-review and AI-assistance platforms relative to MetaBeeAI. The comparison highlights several key axes of differentiation:

- Full-text processing: whether the tool operates directly on PDFs rather than abstracts or metadata.
- User-specified corpus: ability for users to define and manage their own document sets.
- Web literature search: capacity to automatically retrieve documents from online databases or APIs.
- Domain / question agnostic design: flexibility to address user-defined questions across scientific fields without retraining.
- Modular architecture: presence of discrete, replaceable processing stages.
- Source linking: traceable alignment between extracted information and original text segments.
- Benchmarking: inclusion of automated quality evaluation or expert scoring.
- Open-source: whether all source code is available for adaptation and further development.

Collectively, our landscape analysis demonstrates that MetaBeeAI uniquely integrates end-to-end automation with expert oversight, bridging the gap between proprietary summarization tools and fully manual systematic reviews. Its transparent, benchmarked, and adaptable design directly addresses the limitations of existing literature-review platforms, providing a scalable foundation for reproducible, domain-specific evidence synthesis across the biological sciences.

## 3. Materials and Methods

### 3.1. Pipeline overview

MetaBeeAI is a command-line pipeline for performing systematic reviews, designed by and for researchers in the biosciences with the aim of linking the effects of environmental change across levels of biological organization, species, and/or stressor types. The pipeline is demonstrated here within a single, well-curated ecological domain that relies on substantial expert input and domain-specific conventions. While MetaBeeAI was architected with modularity and reuse in mind, its application beyond biosciences has not yet been empirically evaluated. Nevertheless, the underlying design combining expert-in-the-loop validation with a transparent, configurable prompting framework provides a realistic basis for future adaptation to other research domains with comparable evidence structures.

The pipeline (Figure 1) integrates established tools from the scientific Python ecosystem and well-described LLM-based approaches for structured data extraction, alongside a purpose-built graphical user interface for expert annotation and review. The modular design ensures each component: PDF processing and data preparation; text embedding; LLM-based data extraction; and human validation and review; can be independently upgraded as technologies evolve, maintaining long-term sustainability without disrupting existing workflows.

**Figure 1:**
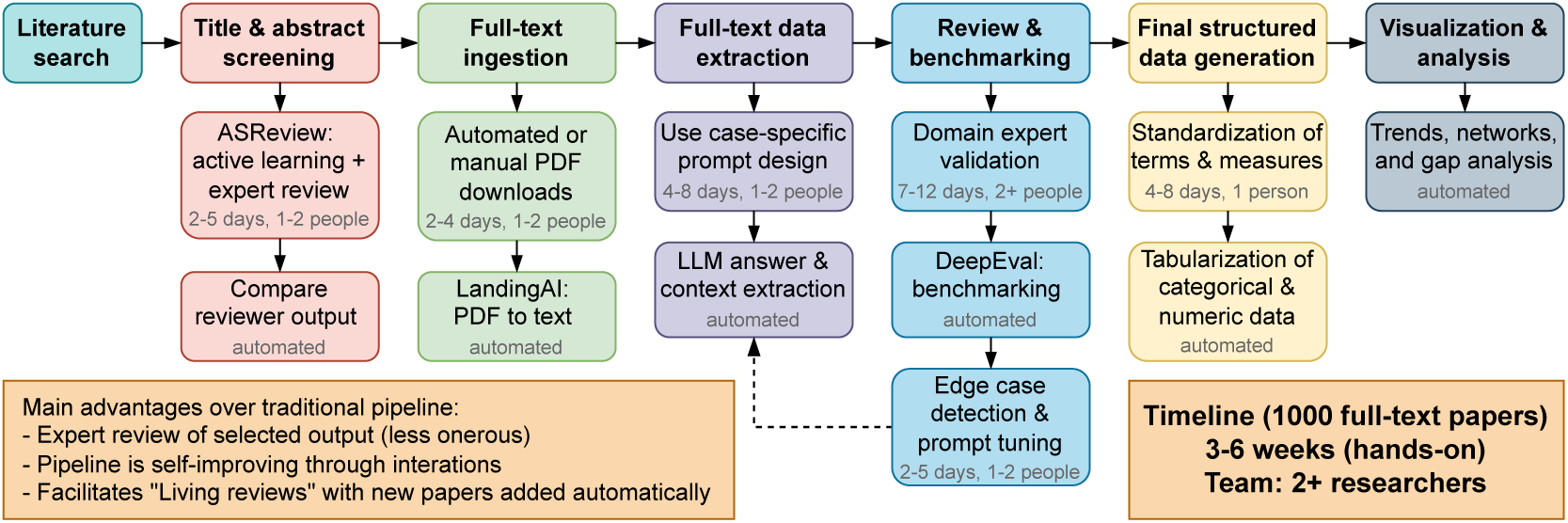
MetaBeeAI is an expert-in-the-loop AI pipeline for enhancing researcher efficiency in the systematic review process. The pipeline begins with a typical search for titles and abstracts of research articles that fit the users’ objectives. Once these have been acquired, the user is guided through each stage of the review, with transparent output at each stage that links any information back to the original paragraph(s) from each article. The ultimate output of the pipeline is a database that can be queried for analyses and synthesis. The suggested timeline is representative of a moderate systematic review including 1000 full-text papers for evidence synthesis.

**Figure 2:**
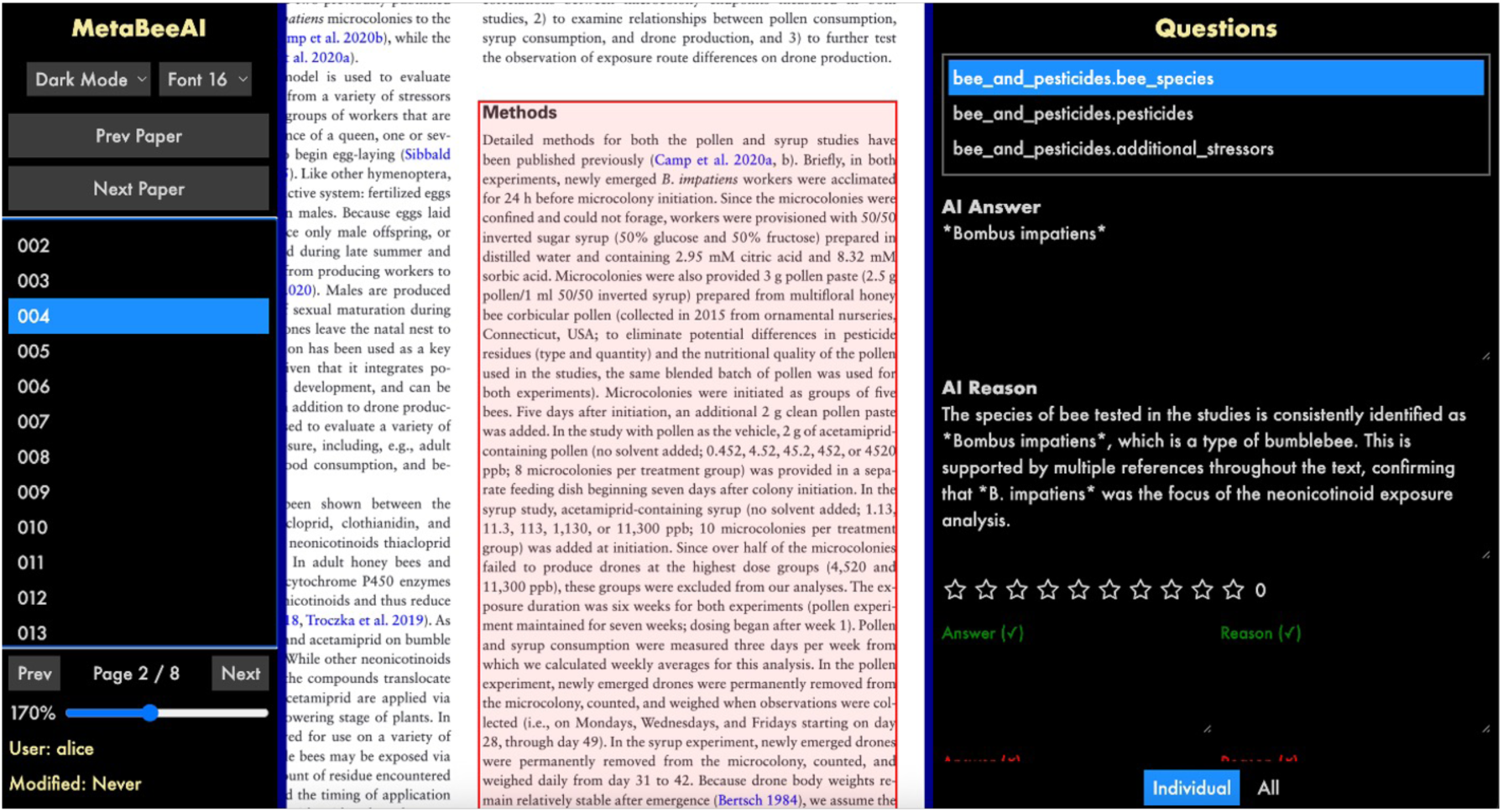
Example screenshot from the review software. The specific paper ID can be selected in the left column, which displays the full text PDF in the centre window. When the user selects a specific question from the right panel, the AI answer and AI reason is printed, and the user can select all or individual relevant text chunks for that question. The user provides a star rating, as well as correct answers and optional LLM answer feedback. Data are automatically stored locally.

MetaBeeAI is distributed as a Python package on PyPI (metabeeai) installable with pip install metabeeai, or users can interact directly with the open-source code base, (GitHub: metabeeai/metabeeai). It provides extensive configuration options via both configuration files and command-line arguments, allowing fine-tuning for a user’s specific use-case and supporting both novice researchers and advanced developers. This is supported by comprehensive documentation (https://metabeeai.readthedocs.io), that includes API references, step-by-step tutorials, and configuration examples.

### 3.2. Pipeline architecture

MetaBeeAI integrates into the systematic review process by combining automated text processing with expert oversight at each stage of the workflow. The process begins with conventional title and abstract screening, in which researchers identify relevant studies from literature searches. To improve efficiency, this step can be assisted by ASReview (developers, 2024), an active-learning screening tool that prioritises abstracts most likely to be relevant based on a small number of user-labelled examples. This allows large bodies of literature to be filtered rapidly while keeping inclusion decisions under direct human control.

For studies that pass screening, full-text PDFs are collected and converted into structured text using Agentic Document Extraction (GitHub: landing-ai/agentic-doc). Each paper is automatically broken into short, paragraph-length sections that retain a direct link to their location in the original document. This step standardises diverse article formats and makes the content easier to query while preserving traceability back to the source text. To reduce costs of the PDF to text conversion, pages containing only irrelevant information (e.g., reference lists) can be manually deleted prior to performing the conversion step.

Information extraction is then carried out in multiple stages. For each research question (for example, species studied, stressors tested, or experimental design), the system first identifies the most relevant text sections within each paper. These sections are then analysed together to extract structured answers, and combined into a single consolidated response per paper. At every stage, the system is designed to return an explicit “no information found” outcome when relevant evidence is absent, reducing over-interpretation. Information querying and extraction is achieved with an LLM, tested here using OpenAI’s GPT-4o and GPT-4o-mini models.

Crucially, all extracted information is presented to expert users through a custom-designed graphical interface that displays the model output alongside the original source text. Researchers can verify, correct, or refine each extracted field, and these refined (“gold”) answers are stored together with the underlying text and extraction settings and can be used to assess the consistency of LLM answers and the convergence of LLM output with expert judgement in the benchmarking stage.

Overall, MetaBeeAI is designed to accelerate evidence synthesis without automating scientific interpretation, providing a structured, auditable bridge between large-scale text processing and expert-led systematic review. A more detailed description of the pipeline architecture, processing steps, and configuration options is provided in Appendix B.

### 3.3. Benchmarking and prompt refinement

The LLM benchmarking system provides evaluation and validation of the LLM-generated answers against human assessment using the DeepEval (ConfidentAI) framework. DeepEval employs LLM-as-a-judge evaluation methods (Zheng et al., 2023), where a user-defined LLM evaluates response quality across multiple dimensions. The system implements a multi-layered evaluation approach that combines various metrics to measure answer quality across dimensions (Table 1). The core evaluation pipeline processes LLM outputs through three primary DeepEval metrics (faithfulness, contextual precision, contextual recall), which compare the LLM output versus the text context (full paper and retrieved text chunks). Additionally, LLM outputs are judged with reference to the expert annotated “gold” answers using two G-Eval metrics (completeness, accuracy). G-Eval (Liu et al., 2023) is a generalized evaluation framework that uses structured, rubric-based prompts to guide LLMs in scoring outputs along defined criteria, improving consistency and alignment with human judgments across tasks such as summarization and question answering. The benchmarking framework incorporates robust batch processing with flexible user-defined model use, with GPT-4o-mini as the default assessment model. The benchmarking system can be used to assess the convergence of LLM output with expert annotations, as well as comparing between experts to demarcate differences in expert annotations of the LLM output.

**Table 1:** Description of benchmarking metrics and the required input data, including LLM answer (output answers from pipeline), gold answer (reviewer-verified benchmarks), retrieval context (relevant text chunks) and context (full paper text).

| Metric | What it measures | Required data |
| --- | --- | --- |
| faithfulness | How well the actual output aligns with both the expected output and the paper context. Detects hallucinations and context violations. | - LLM answer<br>- gold answer<br>- context |
| contextual precision | Evaluates whether the retrieved context contains the specific details needed to answer the question from the gold answer. | - gold answer<br>- retrieval context |
| contextual recall | Completeness of context coverage. Ensures all necessary information was retrieved from the paper. | - gold answer<br>- retrieval context |
| GEval completeness | Coverage of all key points mentioned in the expected output. Assesses if the actual output addresses all important aspects. | - LLM answer<br>- gold answer |
| GEval accuracy | Semantic alignment and information quality. Evaluates if the actual output contains accurate information that aligns with the expected output. | - LLM answer<br>- gold answer |

Following benchmarking with the DeepEval metrics, the pipeline automatically identifies edge cases and generates an analysis of poorly performing LLM responses through comprehensive evaluation score analysis and diagnostic reporting. The system identifies edge cases by analysing evaluation results from DeepEval benchmarking, focusing on responses that received the lowest combined scores across the evaluation metrics. The user can define how many edge cases to include in the final report by changing the relevant setting in the DeepEval framework.

The system then uses these edge cases, their scores on the various metrics, and the automatically generated “reasons” for the scores (generated through the DeepEval framework) to produce a summary report using GPT-4o-mini (or other appropriate model, which can be selected by the user). The output report is a structured, human-readable report that synthesizes edge case findings including the “main issues” that summarize recurring problems for a given question, including specific aspects such as a lack of specificity in responses, inclusion of irrelevant information, omission of key details, and misalignment with expected outputs. The report also includes “recommendations for improving prompt” sections that provide actionable suggestions for prompt modifications (c.f. Section 4.1 for assessment and usage). Each question type analysis includes sample edge cases with specific examples showing the paper, combined scores, input questions, expected output, and problematic LLM responses, enabling researchers to understand failure patterns and develop targeted improvements. Prompts can be updated given these recommendations and the pipeline re-run to quantify improvements. All output, including LLM answers, context (text chunks), benchmarking scores, and reviewer answers, are saved together for further analysis.

### 3.4. Database standardization and analyses

The final step involves the conversion of the LLM answers into a standardized format allowing for analyses. This includes separating the LLM answers from sentences to structured output. The database can then be queried using a noSQL format via JSON files, and additionally can be flattened to a standard CSV file for rapid analyses using statistical software. Standard names for species, locations, experimental perturbations and endpoints, and other variables can be enforced with the use of a dictionary, and units converted to a standard format across fields.

Several analyses can be performed on specific fields selected by the user, including histograms showing the number of papers per field and the distribution of data within categories. Sankey diagrams show relationships between categorical data and highlight topics that are least represented in the data, and can be used to draw inferences about the interconnectivity of the fields.

### 3.5. Case study: bee ecotoxicology

The focal use case for testing the pipeline focused on the topic of nicotinic cholinergic pesticides (e.g., neonicotinoids) and bees. Neonicotinoids are used widely in agriculture, and are often applied prophylactically as seed treatments to crops (Douglas and Tooker, 2015), despite known impacts on beneficial insects like bees (Sgolastra et al., 2020; Hladik et al., 2018). The aim of this systematic review was to highlight the current state of scientific knowledge regarding the diversity of study types including bee species and nicotinic cholinergic pesticides across “domains” of research from molecule to ecosystem, as well as to highlight studies that include pesticides and additional environmental stressors, like temperature extremes and nutritional stress.

We included all primary research papers that had a nicotinic cholinergic pesticide as at least one treatment effect, and included measured results for any species of bees. We included laboratory, semi-field, and field studies. Residue analyses were omitted. An initial corpus of research articles was constructed using the results from literature searches on Scopus and Web of Knowledge with a search term containing two parts, a bee species term and a pesticides term (wherein * denotes a wildcard term):

(bee OR bees OR honey* OR bumble* OR solitary OR stingless OR “wild bees” OR pollinator* OR osmia OR apis OR bombus OR xylocop* OR halictid* OR colletid* OR peponap* OR megachil* OR nomi* OR andren* OR melip*)

AND

(neonicotinoid* OR sulfoximine* OR butenolide* OR cholinergic OR imidacloprid OR clothianidin OR acetamiprid OR thiamethoxam OR dinotefuran OR nitenpyram OR thiacloprid OR acetamiprid OR sulfoxaflor OR spinosyn OR spinosad OR “seed treatment“)

We refined the initial set of papers using ASReview (developers, 2024) removing those that did not fit our inclusion requirements (N = 2 reviewers). We additionally assigned “domains” to each paper, describing the level(s) of biological organization studied in the paper (N = 10 reviewers, with 2 or 3 reviewers per domain). The domains included: molecular (receptor binding, gene expression, metabolism), subindividual (cellular or tissue-level effects), individual (behaviour, fecundity), population (colony-level effects, reproductive rate, survival), and community (field observation of multiple bee species). We included traditional mortality assays (LD50, LC50, etc) as a separate domain, due to the prevalence and traditional importance of these measurements in the field of toxicology (Morris-Schaffer and McCoy, 2021). All abstracts and titles were double-reviewed.

The questions used in the LLM pipeline were:

1. What bee species were experimentally tested in this study?
2. What pesticide(s) were tested in this study? For each, provide the specific dose(s) tested (or field application rates), exposure method(s), and duration of exposure.
3. Were the effects of any additional stressors included in the study (like temperature, parasites or pathogens, other chemicals, or diet and nutrition stress)?

### 3.6. Statistical methods

Statistical analyses on benchmark metrics were conducted separately for each metric, comparing LLM answer convergence with reviewer judgement (LLMv1, LLMv2) and comparing the similarity of reviewer assessment of LLM output. Reviewer star ratings and inter-reviewer start rating differences were analysed using linear mixed-effects models with question type as a fixed effect and paper ID as a random intercept, with significance assessed via likelihood-ratio tests and Bonferroni-corrected post-hoc contrasts. For the benchmarking performance scores, the random paper-level effect explained negligible variance and produced singular fits, so models were simplified to generalised linear models (GLMs) with system, question type, and their interaction as fixed effects. Metrics were expressed as proportions bounded between 0 and 1 and were analysed using Gaussian GLMs applied to logit-transformed scores. Model significance was assessed using type-II ANOVA, and post-hoc pairwise contrasts were performed using estimated marginal means with Tukey adjustment to test differences among systems and question types.

## 4. Results

### 4.1. MetaBeeAI pipeline benchmarking

We selected a subset of 180 papers to test the MetaBeeAI pipeline and obtain domain expert review of the LLM answers (N = 13 reviewers). Of these, 120 papers were double-reviewed so that we could compare LLM versus reviewer judgement, as well as to compare between reviewer judgement of LLM output. The star rating provided a rapid demonstration of differences in reviewer judgement of LLM output across questions, as well as highlighting inter-reviewer differences in assessment (Figure 3). Across all LLM output, answers about bee species received the highest ratings (mean *±* SE = 7.00 *±* 0.27, *n* = 142), followed by pesticides (6.23 *±* 0.21, *n* = 161) and additional stressors (5.56*±*0.33, *n* = 137). After Bonferroni correction, ratings for additional stressors questions remained significantly lower than both bee species questions (*p <* 0.001) and pesticide questions (difference = 0.75, SE = 0.29, *p* = 0.027), whereas the difference between bee species and pesticide questions was not significant (*p* = 0.070). Inter-reviewer agreement, quantified as the absolute difference between two reviewer ratings, did not differ significantly across question types (likelihood ratio test: *χ*^2^(2) = 2.69, *p* = 0.26). Mean inter-reviewer differences were 1.79 *±* 0.30 for bee-species questions (*n* = 56 paired ratings), 2.10 *±* 0.33 for additional-stressors questions (*n* = 51), and 2.46 *±* 0.27 for pesticide questions (*n* = 74), with no significant fixed effects of question type in the mixed-effects model (all *p ≥* 0.10).

**Figure 3:**
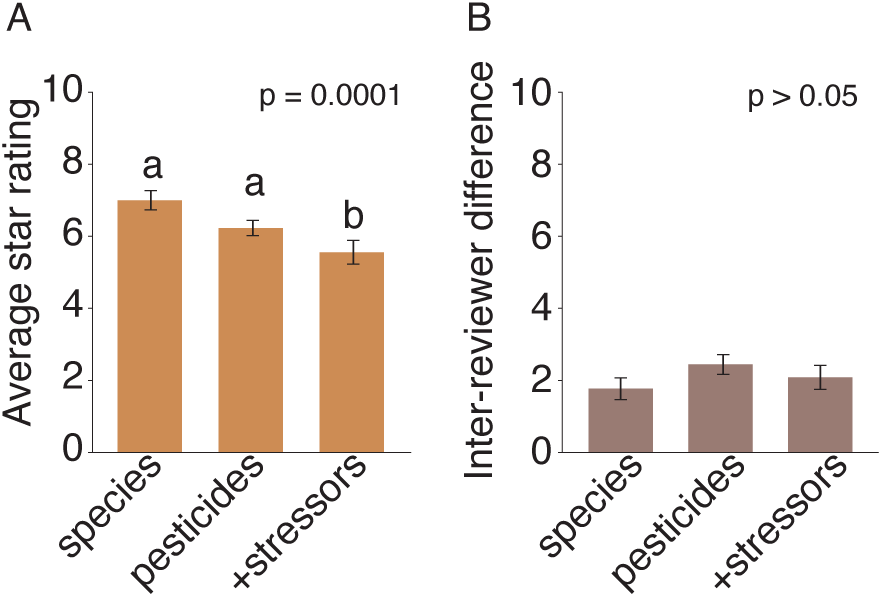
Average reviewer star ratings differed by question type (likelihood ratio test: *χ*^2^(2) = 19.74, *p* = 0.0001) (A), while inter-reviewer agreement, quantified as the absolute difference between two independent reviewer ratings for the same LLM answer, did not differ significantly across question types (likelihood ratio test: *χ*^2^(2) = 2.69, *p* = 0.26) (B).

After performing an initial benchmarking (“LLMv1”), we adapted the prompt pipeline to incorporate the suggestions from the edge case analysis:

1. Emphasise specificity with clearer instructions (e.g., species names).
2. Clarify output requirements (e.g., specify response format and instruct what to omit).
3. Provide examples (e.g., show example responses that are in the correct format).

With these updates, we re-ran the LLM pipeline to obtain a second set of answers (“LLMv2”). We then compared the output from LLMv1, reviewer vs reviewer, and LLMv2 to assess pipeline improvement. Typically, the improvements involved better formatting, a reduction in extra information, and greater answer specificity (Table 2).

**Table 2:** Example answers from LLMv1, reviewers, and LLMv2.

| Question | LLMv1 answer | Reviewer answer | LLMv2 answer |
| --- | --- | --- | --- |
| **bee species** | The species of bees tested were honeybees, specifically *Apis mellifera* and its subspecies *A. m. carnica*. | *Apis mellifera carnica* | 1. *Apis mellifera carnica* |
| **bee species** | The species of bees tested were bumblebees and honey bees. | *Bombus terrestris* and *Apis mellifera* | 1. *Bombus terrestris* 2. *Apis mellifera* |
| **pesticides** | The pesticides used were thiamethoxam and imidacloprid. Exposure was chronic via syrup and pollen at 0, 1.5, 5, 10, and 25 ppb for 18 days. | Bees received only imidacloprid, chronically for 18 days via treated pollen and nectar at 1, 5, 10, and 25 ppb. | 1. Imidacloprid (0, 1, 5, 10, or 25 ppb); oral exposure; 18 days |
| **pesticides** | The pesticides included linalool, hexyl acetate, Z-3-hexenol, (E)-ocimene, terpineol, clothianidin, imidacloprid, and thiacloprid; exposure methods varied; durations not fully available. | Bees were exposed to either thiacloprid or imidacloprid, chronically in sugar solution for 6 weeks (imidacloprid 50 ppb; thiacloprid 4,500 ppb). | 1. Imidacloprid (50 ppb, oral, 6 weeks); 2. Thiacloprid (4,500 ppb, oral via sugar syrup, 6 weeks) |
| **additional stressors** | Yes: DWV infection (1,000 genome copies), nicotine (50 ppm), and 32°C; DWV and nicotine via diet; temperature via thermostatic cabinet; DWV/nicotine until death after emergence; temperature continuous. | DWV infection (1,000 genome copies) and 32°C; DWV via diet; nicotine 50 ppm fed; thermostatic cabinet; DWV/nicotine until death after emergence; temperature continuous. | 1. DWV infection (1,000 genome copies per bee, via diet before pupation, duration n/s); 2. Nicotine (50 ppm in sugar solution at adult stage, duration n/s); 3. Temperature (32°C, duration n/s) |

Across metrics and questions, the LLMv1 scores were typically only slightly lower than the reviewer vs reviewer scores, while the LLMv2 answers demonstrate improvement across metrics (Figure 4). The scores for all metrics except for faithfulness were significantly lower for LLMv1 compared with LLMv2. This effect was driven largely by an improvement in answer specificity and answer formatting, and better handling of missing information, and was especially true for the “additional stressors” question, where LLMv1 would output spurious results when there were no additional stressors mentioned in the paper. In LLMv2, we defined output phrases for missing information, which resulted in higher scores for accuracy and completeness.

**Figure 4:**
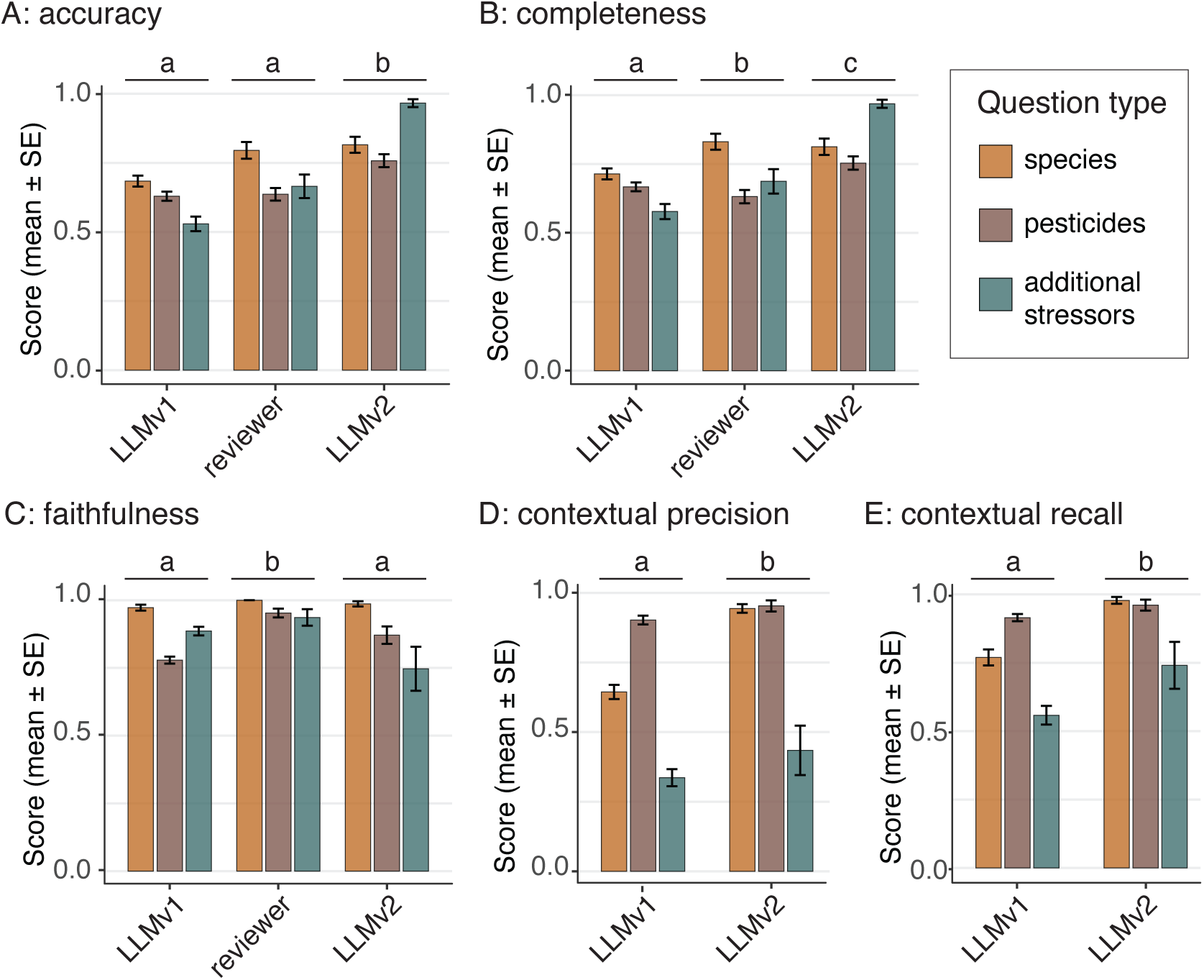
Benchmarking comparing LLMv1 vs reviewer, reviewer vs reviewer, and LLMv2 vs reviewer across accuracy (A, GLM, type: F = 102.3, p < 0.0001; question: F = 12.8, p < 0.0001; type*question: F = 17.9, p < 0.0001), completeness (B, GLM, type: F = 42.2, p < 0.0001; question: F = 5.0, p = 0.0067; type*question: F = 14.3, p < 0.0001), faithfulness (C, GLM, type: F = 26.2, p < 0.0001; question: F = 100.9, p < 0.0001; type*question: F = 12.4, p < 0.0001), contextual precision (D, GLM, type: F = 66.7, p < 0.0001; question: F = 196.3, p < 0.0001; type*question: F = 12.4, p < 0.0001) and contextual Recall (E, GLM, type: F =34.4, p < 0.0001; question: F = 57.0, p < 0.0001; type*question: F = 3.4, p = 0.035) metrics. Letters denote significance between sources (LLMv1, reviewer vs reviewer, LLMv2). Description of metrics can be found in Table 1.

The total cost of benchmarking across the three evaluation runs (LLMv1, LLMv2, reviewer vs reviewer, 194 papers with 3 questions per paper, total of 1,746 evaluations) was US $21.43. The majority of the cost going towards the Faithfulness (43%), Contextual Precision (20.9%), and Contextual Recall (15.7%) metrics, which require processing of the the full text and text chunks from each paper. These metrics required the use of GPT-4o given the higher processing demand. The two G-Eval metrics were cheaper, representing a combined 17.7% of the total cost, and could be run using the lighter GPT-4o-mini model.

### 4.2. Abstract and Title Screening: bees and pesticides literature

The literature searches identified 4,509 papers. This was reduced to 1,053 papers after filtering titles and abstracts using the inclusion criteria. We applied domain labels to each paper, based on the focus of the experiments or observations in each paper (Figure 5). A single paper could have one or multiple domains, and some papers (186) were not given a domain label. There was an average of 83% agreement between reviewers for domain labels. The 186 papers that had not been allocated to a domain were re-reviewed, and 103 of these were removed due to mislabelling at the initial filtering stage. A total of 924 papers were available as full-text, proceeding through the remainder of the pipeline, including PDF to text conversion and the LLM data extraction system.

**Figure 5:**
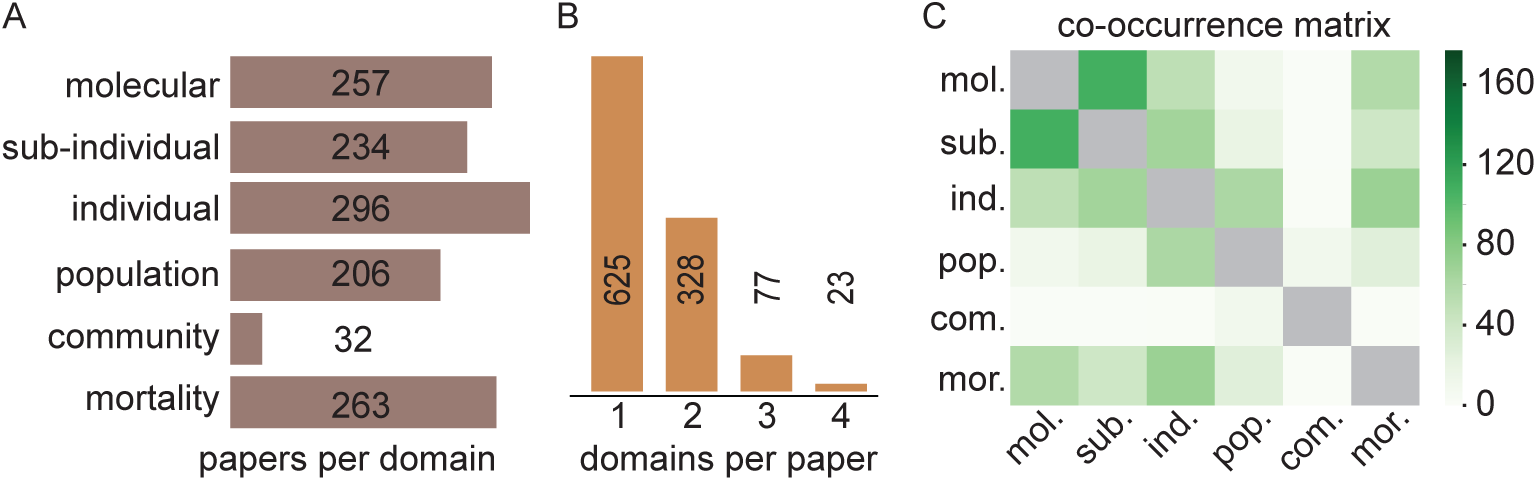
Paper topic distribution across levels of biological organization and domain (A), the number of domains addressed within each paper (B), and the co-occurrence of domains within papers (C).

### 4.3. Case study: bees and pesticides literature review

Across all the papers in the database, there were 339 unique bee species and 14 unique nicotinic cholinergic pesticides, amounting to 438 unique bee-pesticide combinations. When filtering out studies that included pollinator community surveys, there were 130 unique bee species from 44 genera (Figure 6). *Apis* (honeybees) were studied in 672 papers, followed by *Bombus* (bumblebees) in 223 papers, *Osmia* (mason bees) in 83 papers, *Melipona* (stingless bees) in 33 papers, and *Megachile* (leafcutter bees) in 26 papers.

**Figure 6:**
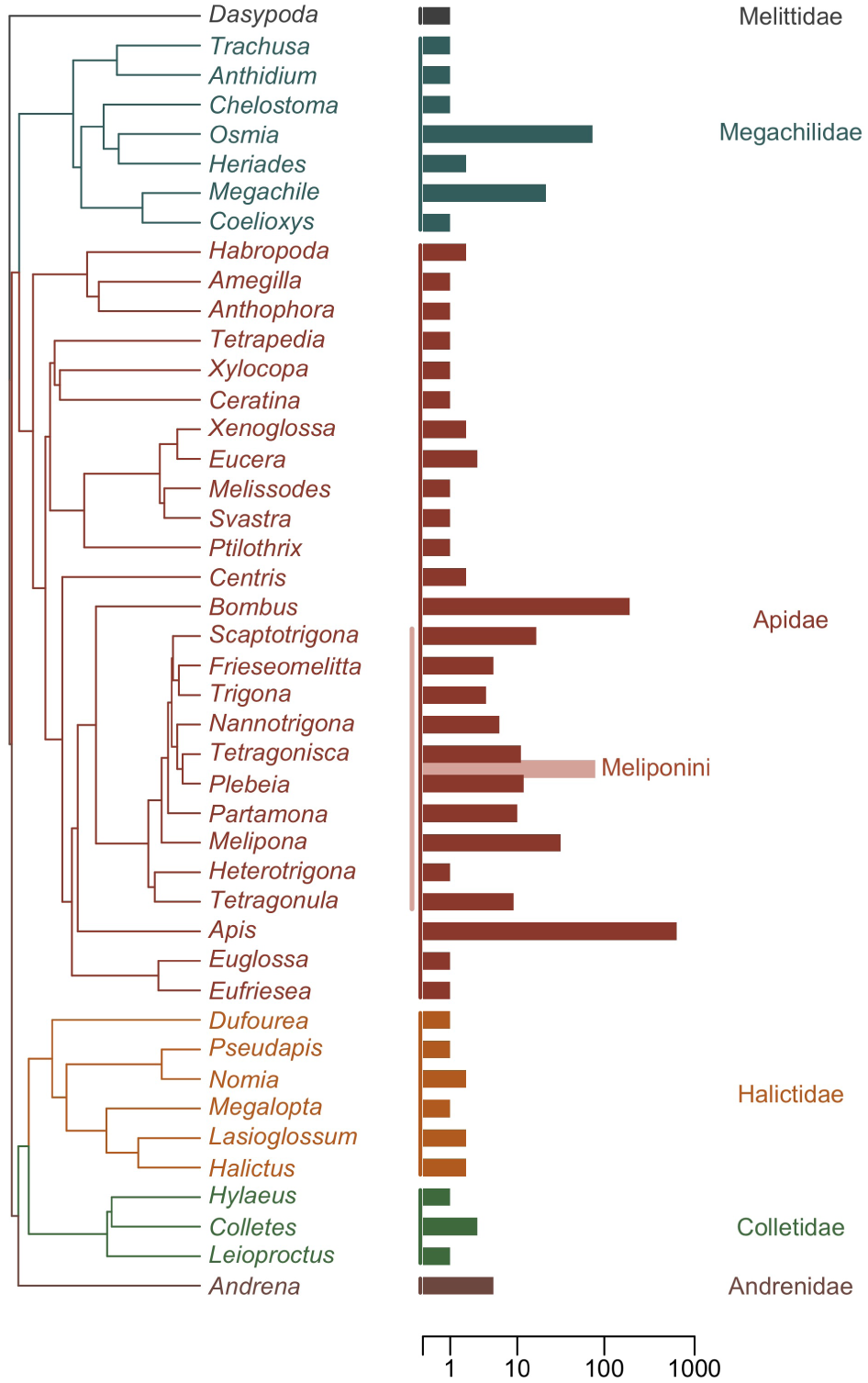
Phylogenetic tree adapted from Henríquez-Piskulich et al. (2024) showing the genera included in the database of papers testing the effects of nicotinic cholinergic pesticides on bees (left), and the number of papers per genus (right). Community survey studies were removed and genera only represented by such studies have been excluded. Vertical coloured lines at x = 0 denote families, with family names displayed on the right of the bars. Genera belonging to the tribe Meliponini (stingless bees) are in addition marked with a pink vertical line and the corresponding papers (n = 79) are summarized by a pink bar. To improve readability, the x-axis has been compressed. Honeybees (*Apis*, n = 637) and bumblebees (*Bombus*, n = 192) represent the two most studied genera. In total, 842 of the papers (excluding community surveys) include at least one species from the monophyletic clade consisting of these three eusocial groups (Meliponini, *Apis* and *Bombus*), whereas 57 papers do not.

We explored the relationships between bee species, pesticides, and additional stressors (Figure 7). Here, “study” refers to the set of experiments within a paper for a given bee and pesticide combination, so a single paper can have multiple studies (i.e., multiple species and/or pesticides, or the same species included in more than one experiment). Unsurprisingly, *Apis mellifera* (honeybee) represented by far the largest number of studies across the main neonicotinoid pesticides (total: 813 studies; imidacloprid: 266 studies; thiamethoxam: 155 studies; clothianidin: 138 studies), followed by *Bombus terrestris* (bumblebee, total: 176 studies; imidacloprid: 55 studies; thiamethoxam: 38 studies). When considering all pesticide studies, *Apis cerana* was the third most studied (75 studies) followed by *Bombus impatiens* (71 studies), *Osmia bicornis* (67 studies), *Megachile rotundata* (31 studies), *Melipona quadrifasciata* (24 studies), *Osmia lignaria* (24 studies), *Apis florea* (18 studies), and *Scaptotrigona postica* (13 studies).

**Figure 7:**
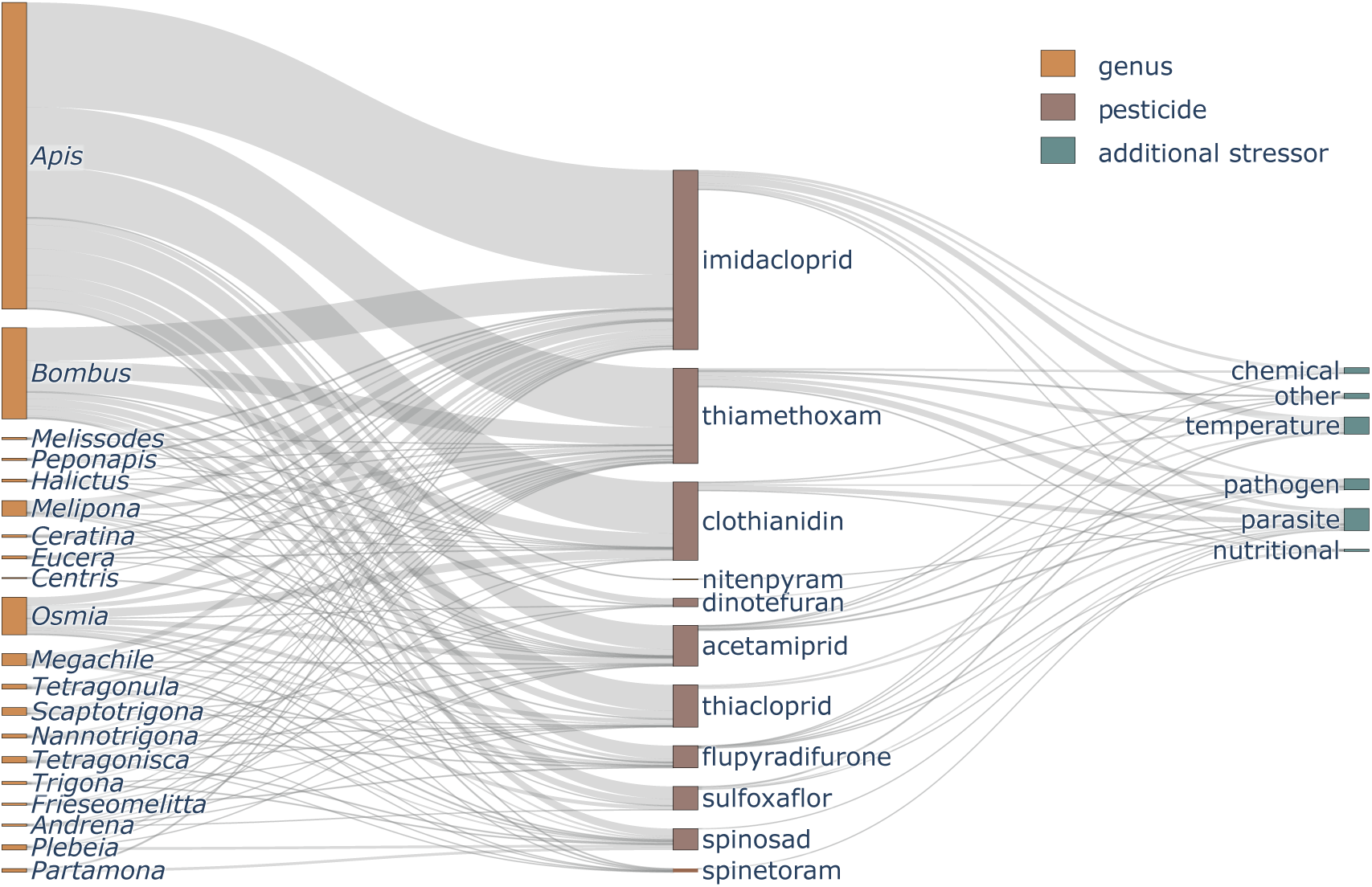
Sankey diagram showing the co-occurrence of bee genera with pesticide types (filtered with min. 3 studies per relationship) and pesticides with additional stressor categories. Thicker lines represent a larger number of studies.

Secondary stressors were included in 182 studies across 106 papers, comprised of 6 broad stressor groups: parasites (55 studies), temperature (45 studies), pathogens (32 studies), chemical stressors (31 studies), 14 “other” stressors (e.g., UV-light), and 5 papers that included nutritional stress.

Running the MetaBeeAI pipeline on 924 papers with three extraction questions incurred a total cost of US$443.52. This included PDF-to-text conversion using Agentic Document Extraction (US$277.20) at US$0.03 per page, and the multi-step large language model (LLM) information extraction pipeline (US$166.32). The latter corresponds to a cost of US$0.06 per question, applied to three questions across 924 papers, using GPT-4o for identification and relevance scoring of candidate text chunks as well as answer extraction from relevant chunks. The total time required to convert the PDFs to text was approximately 16 hours, given the rate limitations from LandingAI. The LLM pipeline required 31 hours to process all papers. All processing was performed on a standard laptop with a stable internet connection, as all interaction with LLMs occurs via API. Overall costs (financial and temporal) could be reduced by substituting GPT-4o-mini for one or both stages of the LLM pipeline, although this may come at the expense of extraction accuracy. The impact of model choice on answer quality, as well as performance of alternative LLMs, remains to be systematically evaluated.

## 5. Discussion

### 5.1. Performance in a bee ecotoxicology case study

We evaluated the MetaBeeAI pipeline on 924 research articles examining the effects of nicotinic cholinergic pesticides across molecular, behavioural, colony, and field-level endpoints. This corpus provided a demanding test due to its heterogeneity in terminology, experimental design, and reporting standards. Evaluation focused on expert judgement of LLM-generated extractions rather than comparison against an independently constructed benchmark dataset.

The pipeline performed reliably for factual extraction tasks such as identifying compounds and focal taxa, but showed more variable performance for tasks requiring contextual interpretation. Common failure modes included incomplete mapping between commercial formulations and active ingredients, ambiguous assignment of pesticide classes when information was implicit, and inconsistent normalization of species names. These issues were identified through expert review of model outputs rather than automated scoring.

Iterative refinements informed by workshop discussions with domain experts – particularly updates to prompt dictionaries, chemical mappings, and ontology files – improved consistency and recall in subsequent runs. This process highlights the role of expert-in-the-loop validation in diagnosing systematic error modes and incrementally improving extraction quality. The case study therefore serves primarily as a diagnostic and developmental evaluation, rather than a definitive benchmark of extraction accuracy.

### 5.2. Generalisability and broader applications

MetaBeeAI is designed as a modular framework rather than a domain-specific solution. In principle, its hierarchical prompting, expert-in-the-loop validation, and benchmarking-by-review structure could be adapted to other scientific domains by modifying prompt files, dictionaries, and schemas. However, this generalisability has not yet been empirically tested beyond bee ecotoxicology.

The framework is intended to occupy a middle ground between generic AI literature assistants and fully manual systematic reviews, providing structured outputs with explicit links to source text. An additional by-product of expert validation is the creation of curated datasets of model outputs and reviewer judgements, which may support future model comparison or fine-tuning. Whether this approach scales effectively to other fields, publication formats, or evidence types remains an open question for future work.

### 5.3. Comparison with other LLM pipelines

MetaBeeAI differs primarily in its emphasis on full-text extraction, explicit linkage between outputs and source passages, and iterative expert validation. Its modular architecture allows individual components—such as document parsing, relevance filtering, or schema design—to be updated independently, reducing coupling to specific models or tools. This design prioritizes transparency and auditability rather than fully automated synthesis.

MetaBeeAI is therefore best viewed as complementary to general-purpose literature assistants. While other systems facilitate rapid exploration, MetaBeeAI is intended to support structured evidence mapping and data generation under expert oversight, albeit with higher human involvement and currently limited validation outside the presented case study.

### 5.4. Strengths, limitations, and future directions

Key strengths of MetaBeeAI include its modular design, transparent linkage between extracted statements and source text, and reliance on expert review to assess and refine LLM outputs. This structure helps mitigate known risks such as hallucination and context misinterpretation (Bang et al., 2023).

Several limitations remain. Extraction quality depends on upstream PDF-to-text conversion, which can introduce artifacts. Numerical data from tables and figures were not reliably extracted and were excluded from downstream use. Harmonization of terminology, units, and taxonomies currently requires manual curation, constraining scalability. As with other LLM-based systems, reproducibility is affected by stochastic inference and ongoing model updates (He, 2025), though MetaBeeAI preserves traceability to support re-evaluation as models change.

A further limitation concerns potential cognitive bias during expert review. Presenting model-generated answers prior to human assessment may induce anchoring effects (Tversky and Kahneman, 1974), influencing reviewer judgement. This risk has not been formally tested here. Future work should evaluate alternative review protocols, such as blinded assessment or independent expert annotation prior to exposure to model outputs.

Future development could include tighter integration with domain ontologies, improved table-parsing models for numerical extraction, and community-curated evaluation datasets. Advances in multimodal models may eventually allow incorporation of figures and images, though this remains untested within the current framework.

### 5.5. Conclusion

MetaBeeAI provides a transparent, expert-in-the-loop framework for structured extraction of information from scientific literature. Applied here to bee ecotoxicology, it demonstrates how modular AI tools combined with human judgement can support scalable evidence synthesis while retaining traceability and oversight. Although the framework is designed to be adaptable, its performance and utility in other domains have not yet been evaluated. As scientific literature continues to grow in volume and complexity, approaches such as MetaBeeAI may help bridge the gap between automated text processing and the standards required for systematic, reproducible synthesis, provided their limitations and assumptions are made explicit.

## 7. Author contributions

R. H. Parkinson led the conceptualization, methodology design, software development, validation, formal analysis, investigation, resources, data curation, visualization, supervision, project administration, writing, and funding acquisition. H. Cerbone contributed to methodology, software development, validation, formal analysis, investigation, resources, data curation, writing, supervision, and project administration. S. Cao contributed to conceptualization, methodology, software development, resources, writing, and funding acquisition. M. Mieskolainen contributed to the conceptualization and software development and validation. A. D. Wilson contributed to software development and validation. S. Roberts contributed to conceptualization, supervision, writing, and funding acquisition.

S. Albacete, E. B. Armstrong, A. Brown, C. Botías, A. K. Jones, H. Siviter, F. Sgolastra, D. A. Stanley, E. A. Straw, A. J. Hayward, L. Herbertsson, N. Nagloo, E. Rigosi, E. Nicholls, R. Tadei, and K. Walter contributed to methodology, investigation and manuscript review. L. Herbertsson also contributed to visualization. C. Bass contributed to investigation, writing, and supervision. L. Straub contributed to manuscript review. R. K. Daniels and H. F. Stevance contributed to methodology, validation, and manuscript review. B. Lambert contributed to writing and funding acquisition.

All authors reviewed and approved the final version of the manuscript.

## 8. Declaration of generative AI use

During the preparation of this work, the authors used OpenAI’s GPT-4.0-mini, GPT-4.0, and GPT-5 models to assist in refining text and summarising aspects of the pipeline methodology. All content generated with these tools was reviewed, verified, and edited by the authors, who take full responsibility for the final text and its scientific accuracy.

## 9. Data availability

The data used in the generation of this manuscript are available on FigShare (https://doi.org/10.6084/m9.figshare.31209130). All code are available on GitHub (https://github.com/MetaBeeAI/MetaBeeAI).

## Appendix A. Example YAML Schema for Pesticide Extraction Task

pesticides:

question: “What pesticide(s) were tested in this study? For each, provide the specific dose(s) tested (or field application rates), exposure method(s) and duration of exposure.”

instructions:

- “Report pesticides that bees were exposed to in the study (including field studies, controlled experiments, etc.).”
- “Use the chemical name (not trade name or formulation) unless no other name is given.”
- “Include doses/concentrations if explicitly reported, or field application rates if explicitly reported.”
- “Include exposure method if explicitly reported (oral, contact, topical, field exposure, etc.).”
- “Include exposure duration if explicitly reported.”
- “List each pesticide separately.”
- “Do not add any interpretation, context, or description of results.”
- “Do not infer missing information—only report what is explicitly stated.”
- “If only the pesticide name is available, report just the name.” output_format: “List of pesticides with: chemical name, and any available details (dose, exposure method, duration)” example_output:
- “1. Imidacloprid: 10 and 100 ppb, oral exposure, 7 days; 2. Thiamethoxam: 5 and 25 ppb, contact exposure, acute”
- “1. Clothianidin: field exposure via treated oilseed rape at 10 g per kg of seed; 2. Beta-cyfluthrin: field exposure via treated seeds”
- “1. Clothianidin; 2. Indoxacarb” bad_example_output:
- “In the study, the researchers tested imidacloprid (Admire) and sufoxaflor (Closer) by feeding bees the pesticides over 10 days.”
- “The study tested several pesticides including neonicotinoids which are harmful to bees. They used different doses and exposed the bees for some time. The

pesticides were applied in various ways to see how they affected the bees.” no_info_response: “No pesticides were tested in this study” max_chunks: 5

**Table A.3:** Comparison of literature-review / LLM pipelines and capabilities. All metrics are based on information available on the listed tool’s homepage (Lála et al., 2023; Paper Digest, 2025; Elicit, 2025; Marshall et al., 2017; Inc., 2025; Consensus, 2024; LLC, 2025; DistillerSR Inc., 2025; Veritas Health Innovation, 2025; Inc, 2025; ReadCube, 2025; Tilda Technologies Inc., 2025; Paper Digest, 2025; Documind, 2025; Anara Labs Inc, 2025; Elsevier, 2025; Allen Institute for AI, 2024).

| Pipeline | Full-text processing | User-specified corpus | Web literature search | Domain & question agnostic | Modular architecture | Source linking | Bench-marking | Open-source |
| --- | --- | --- | --- | --- | --- | --- | --- | --- |
| MetaBeeAI | yes | yes | no | yes | yes | yes | yes | yes |
| PaperQA2 | yes | yes | yes | yes | yes | yes | yes | yes |
| Elicit | yes | partial | yes | yes | no | yes | yes | no |
| RobotReviewer | yes | yes | no | no | yes | yes | yes | yes |
| SciSpace | yes | partial | yes | yes | no | yes | yes | no |
| Consensus | yes | no | yes | yes | no | yes | no | no |
| Scite.ai | yes | yes | yes | yes | no | yes | no | no |
| DistillerSR | yes | yes | yes | yes | yes | yes | no | no |
| Covidence | yes | yes | yes | yes | yes | yes | no | no |
| Perplexity (Deep Research) | partial | no | yes | yes | no | yes | yes | no |
| ReadCube / Papers AI | yes | yes | yes | yes | no | yes | no | no |
| Humata | yes | yes | no | no | no | yes | no | no |
| PaperDigest | yes | yes | yes | yes | no | yes | no | no |
| Documind | yes | no | no | yes | yes | yes | no | no |
| Anara | yes | yes | yes | yes | no | yes | no | no |
| Scopus AI | yes | no | partial | yes | no | yes | no | no |
| Asta | yes | partial | yes | yes | no | yes | no | no |

## Appendix B. MetaBeeAI pipeline software methods

### Appendix B.1. Abstract and title screening

To ensure the inclusion of articles with a relevant scope, users review the titles and abstracts of the papers found during an initial search. We recommend compiling the results from two or more sources to ensure appropriate coverage. Abstract screening is performed in ASReview v1.6.2 (van de Schoot et al. (2021), GitHub: asreview/asreview), which uses active learning to sort abstracts based on relevance. The user establishes relevant papers for inclusion and exclusion with a small subset of examples, following which the model sorts the corpus of abstracts from best to worst. The graphical user interface is easy to navigate, and enhances the speed at which the researcher can evaluate a large number of abstracts.

Once the corpus of relevant papers has been established, reviewers need to download the full text PDFs. This can be done automatically using a free-to-use tool like Unpaywall (unpaywall.org) or similar for open access papers.

### Appendix B.2. Conversion of PDFs to structured text

Next, Agentic Document Extraction (ADE) from LandingAI v1 (GitHub: https://github.com/landingai/agentic-doc) is used to convert full-text documents (Portable Document Format, PDF) to a structured JSON (JavaScript Object Notation) format, a text-based format for data interchange that is human-readable and easy for machines to parse. Prior to document conversion, PDFs are separated into one or two-page PDFs for processing due to LandingAI API restrictions. Pages containing only reference lists can be manually removed at this point to reduce cost. The MetaBeeAI pipeline processes each paper in the database, and automatically stitches together the resulting JSON files with the appropriate page reference for the original PDF. Duplicate text chunks are removed (accounting for the page overlaps). We include only the conversion of the main text and figure captions for subsequent processing (header, marginalia, and figures are automatically removed), as the numerical output from ADE which translates images (e.g., figures) to text descriptions was not sufficiently reliable in this version. We anticipate that the functionality that this tool promises will be highly valuable in future iterations of the model for directly extracting data from figures. The result from this step is a series of structured JSON files that are presented as “text chunks” of approximately one paragraph long, plus bounding box coordinates that relate the text chunk back to the location of the text in the original PDF.

### Appendix B.3. Multi-pass extraction and prompt chaining

The MetaBeeAI LLM pipeline implements a multi-pass extraction system for querying structured text chunks and extracting domain-specific information from research papers. The pipeline consists of three main processing stages: chunk relevance filtering, individual chunk querying, and answer synthesis. The system begins by loading question-specific configurations from a YAML file (a human-readable data serialization language that is often used for writing configuration files), which is the main point of interaction of the user with the pipeline (Figure A.1). This file defines the prompt question, in addition to:

- instructions: detailed guidance for information extraction.
- output_format: structured response template.
- example_output: positive examples demonstrating desired responses.
- bad_example_output: negative examples showing patterns or content to avoid.
- no_info_response: the specific wording to use if the desired information is not present in the text.
- max_chunks: the number of top-ranked chunks to select for answer generation.

The system uses a dedicated relevance model (tested with GPT-4o-mini and GPT-4o) to evaluate text chunks against specific questions through a two-stage filtering process. First, rule-based pre-filtering removes obviously irrelevant content (headers, metadata, logos, journal info, DOI, authors, etc.) using keyword matching. Second, LLM-based relevance selection uses a single API (Application Programming Interface) call with a structured prompt containing the question, instructions, output format, good examples, and bad examples from the question configuration. The LLM iterates over all chunks filtered as above and returns an order of relevance over the chunks, selecting the top N most relevant (configurable via max_chunks parameter in the YAML configuration). This approach eliminates unnecessary LLM calls by filtering out irrelevant content early while ensuring question-specific relevance through example-driven prompting.

Based on empirical testing and cost-performance analysis in terms of time, we recommend max_chunks values of 3-6 chunks: lower values (3-4) for focused questions like “future research directions” that typically appear in specific sections, and higher values (5-6) for comprehensive questions like “experimental methodology” that may span multiple paragraphs. This range balances comprehensive coverage with processing efficiency, as each additional chunk increases both answer generation costs and synthesis complexity, while values below three may miss relevant information and values above six show diminishing returns with increasing costs.

The selected relevant chunks are then processed in parallel batches to optimize API usage and processing speed (tested with GPT-4o-mini and GPT-4o). For each chunk, the system constructs a structured prompt containing the question, instructions, output format, good examples, bad examples, text content (the actual chunk text), and quality guidelines emphasizing conciseness, format compliance, and avoiding speculation. Each chunk query returns a structured response containing the extracted answer and reasoning, formatted to ensure consistent output structure.

The final stage consolidates individual chunk responses into a coherent answer through an LLM powered self-reflection process. The synthesis prompt includes: all relevant chunk texts and their individual answers, as well as the various guidelines from the user prompt (instructions, good and bad examples, output structure), and conflict resolution instructions for handling contradictory information across chunks. The reflection stage can return an INSUFFICIENT_INFO flag when available information is inadequate, which is then replaced with question-specific no_info_response text. Additionally, if no relevant chunks are found during the initial filtering stage, the system immediately returns the no_info_response text without proceeding to answer generation or reflection.

The system implements several performance optimizations, including parallel processing (answer generation occurs in configurable batch sizes), adaptive batch sizing (batch sizes adjust based on available chunks and API rate limits), and asynchronous processing (all LLM interactions use async/await patterns for optimal throughput). The multi-pass architecture ensures high-quality information extraction by combining automated relevance filtering with example-driven prompting and iterative refinement, which can leverage different tiers of models for specific steps to enhance output quality while optimizing API costs.

### Appendix B.4. Expert-in-the-loop review interface

We have developed software to function as a comprehensive manual validation system for assessing the accuracy and quality of LLM-generated answers against original research papers and the specific portions of the paper used in generating the answers. The system consists of two complementary tools: an automated PDF annotation system and an interactive graphical user interface (GUI) for detailed manual review of the answers with the original full text PDFs.

The annotation system creates visual overlays on original PDF documents to highlight the specific text regions used by the LLM pipeline for answer generation (Figure 2). This enables reviewers to quickly locate and examine the source material underlying each extracted answer. In drawing the bounding boxes, the annotation software leverages the bounding box output by the PDF to text conversion step, the relevant chunk IDs related to each question, and the original full text PDF.

The PyQt5-based desktop application provides a comprehensive interface for systematic manual review and validation of LLM-generated answers. The user can easily move through different questions and papers, and can select individual text chunks for further investigation. The user can assign a star rating (0 to 10) for each LLM answer, and provide correct answers and reasoning alongside any criticism of the LLM answer or reasoning. The positive answers and reasons provide dual functions for use in pipeline benchmarking (section 3.3) as well as the construction of a domain-specific database of benchmarking questions that could serve for model fine-tuning for those building custom models.

The integrated review system enables systematic human evaluation of LLM performance while maintaining direct visual connection to source materials, supporting both individual answer validation and large-scale quality assessment.

